## Supplemental Tables and Figures for "Dynamic landscape of the intracellular termini of acid-sensing ion channel 1a"

**Figure 1 – figure supplement 1**

DNA sequence for ASIC1a from rat used in this study. Conservative mutations to lessen alternative initiation highlighted in cyan and cysteine to serine mutations highlighted in yellow. Amino acid translation is above in red.

```
M  E  L  K  T  E  E  E  E  V  G  G  V  Q  P  V  S  L  Q  A
ATGGAACTCAAAACCGAGGAGGAGGAGGTCGGTGGTGTCCAGCCGGTCAGCCTCCAGGCT
         10        20        30        40        50

F  A  S  S  S  T  L  H  G  L  A  H  I  F  S  Y  E  R  L  S
TTCGCCAGCAGCTCCACCCTTCATGGTCTCGCCCACATCTTCTCCTATGAGCGGCTGTCT
         70        80        90        100       110

L  K  R  A  L  W  A  L  C  F  L  G  S  L  A  V  L  L  C  V
CTGAAGCGGGCACTGTGGGCCCTGTGCTTCCTGGGTTCGCTGGCCGTCCTGCTGTGTGTG
         130       140       150       160       170

C  T  E  R  V  Q  Y  Y  F  C  Y  H  H  V  T  K  L  D  E  V
TGCACTGAGCGTGTGCAGTACTACTTCTGCTATCACCACGTCACCAAGCTTGACGAAGTG
         190       200       210       220       230

A  A  S  Q  L  T  F  P  A  V  T  L  C  N  L  N  E  F  R  F
GCTGCCTCCCAGCTCACCTTCCCTGCTGTCACACTGTGCAATCTCAATGAGTTCCGCTTT
         250       260       270       280       290

S  Q  V  S  K  N  D  L  Y  H  A  G  E  L  L  A  L  L  N  N
AGCCAAGTCTCCAAGAATGACCTGTACCATGCTGGGGAGCTGCTGGCCCTGCTCAACAAC
         310       320       330       340       350

R  Y  E  I  P  D  T  Q  M  A  D  E  K  Q  L  E  I  L  Q  D
AGGTATGAGATCCCGGACACACAGATGGCTGATGAAAAGCAGCTAGAGATATTGCAGGAC
         370       380       390       400       410

K  A  N  F  R  S  F  K  P  K  P  F  N  M  R  E  F  Y  D  R
AAGGCCAACTTCCGGAGCTTCAAGCCCAAGCCCTTCAACATGCGTGAATTCTACGACAGA
         430       440       450       460       470

A  G  H  D  I  R  D  M  L  L  S  C  H  F  R  G  E  A  C  S
GCGGGGCACGATATTCGAGACATGCTGCTCTCGTGCCACTTCCGTGGGGAGGCCTGCAGC
         490       500       510       520       530

A  E  D  F  K  V  V  F  T  R  Y  G  K  C  Y  T  F  N  S  G
GCTGAAGATTTCAAAGTGGTCTTCACTCGGTATGGGAAGTGTTACACATTCAACTCGGGC
         550       560       570       580       590
```

```
Q  D  G  R  P  R  L  K  T  M  K  G  G  T  G  N  G  L  E  I
CAAGATGGGCGGCCACGGCTGAAGACCATGAAAGGTGGGACTGGCAATGGCCTGGAGATC
         610       620       630       640       650

M  L  D  I  Q  Q  D  E  Y  L  P  V  W  G  E  T  D  E  T  S
ATGCTGGACATTCAGCAAGATGAATATTTGCCTGTGTGGGGAGAGACCGACGAGACATCC
         670       680       690       700       710

F  E  A  G  I  K  V  Q  I  H  S  Q  D  E  P  P  F  I  D  Q
TTCGAAGCAGGCATCAAAGTGCAGATCCACAGTCAGGATGAACCCCCTTTCATCGACCAG
         730       740       750       760       770

L  G  F  G  V  A  P  G  F  Q  T  F  V  S  C  Q  E  Q  R  L
CTGGGCTTTGGCGTGGCTCCAGGTTTCCAGACGTTTGTGTCTTGCCAGGAGCAGAGGCTC
         790       800       810       820       830

I  Y  L  P  S  P  W  G  T  C  N  A  V  T  M  D  S  D  F  F
ATCTACCTGCCCTCACCCTGGGGCACCTGCAATGCTGTTACCATGGACTCGGATTTCTTC
         850       860       870       880       890

D  S  Y  S  I  T  A  C  R  I  D  C  E  T  R  Y  L  V  E  N
GACTCCTACAGCATCACTGCCTGCCGGATTGATTGCGAGACGCGTTACCTGGTGGAGAAC
         910       920       930       940       950

C  N  C  R  M  V  H  M  P  G  D  A  P  Y  C  T  P  E  Q  Y
TGCAACTGCCGTATGGTGCACATGCCAGGGGACGCCCCATACTGCACTCCAGAGCAGTAC
         970       980       990       1000      1010

K  E  C  A  D  P  A  L  D  F  L  V  E  K  D  Q  E  Y  C  V
AAGGAGTGTGCAGATCCTGCCCTGGACTTCCTAGTGGAGAAAGACCAGGAATACTGCGTG
         1030      1040      1050      1060      1070

C  E  M  P  C  N  L  T  R  Y  G  K  E  L  S  M  V  K  I  P
TGTGAGATGCCTTGCAACCTGACCCGCTACGGCAAGGAGCTGTCCATGGTCAAGATCCCA
         1090      1100      1110      1120      1130

S  K  A  S  A  K  Y  L  A  K  K  F  N  K  S  E  Q  Y  I  G
AGCAAAGCCTCCGCCAAGTACCTGGCCAAGAAGTTCAACAAATCGGAGCAGTACATAGGG
         1150      1160      1170      1180      1190

E  N  I  L  V  L  D  I  F  F  E  V  L  N  Y  E  T  I  E  Q
GAGAACATTCTGGTGCTGGACATTTTCTTTGAAGTCCTCAACTATGAGACCATCGAGCAG
         1210      1220      1230      1240      1250

K  K  A  Y  E  I  A  G  L  L  G  D  I  G  G  Q  M  G  L  F
```

```
AAAAAGGCCTATGAGATCGCAGGGCTGTTGGGTGACATCGGGGGCCAGATGGGGTTGTTC
         1270      1280      1290      1300      1310

I  G  A  S  I  L  T  V  L  E  L  F  D  Y  A  Y  E  V  I  K
ATCGGTGCCAGCATCCTCACCGTGCTGGAACTCTTTGACTATGCCTACGAGGTCATTAAG
         1330      1340      1350      1360      1370

H  R  L  S  R  R  G  K  C  Q  K  E  A  K  R  S  S  A  D  K
CACAGGCTGTCCAGACGTGGAAAGTGCCAGAAGGAGGCTAAGAGGAGCAGCGCAGACAAG
         1390      1400      1410      1420      1430

G  V  A  L  S  L  D  D  V  K  R  H  N  P  S  E  S  L  R  G
GGCGTGGCGCTCAGCCTGGATGACGTCAAAAGACACAATCCCTCCGAGAGCCTCCGAGGA
         1450      1460      1470      1480      1490

H  P  A  G  M  T  Y  A  A  N  I  L  P  H  H  P  A  R  G  T
CATCCTGCCGGGATGACGTACGCTGCCAACATCCTACCTCACCATCCCGCTCGAGGCACG
         1510      1520      1530      1540      1550

F  E  D  F  T  S
TTTGAGGACTTTACCTCC
         1570
```

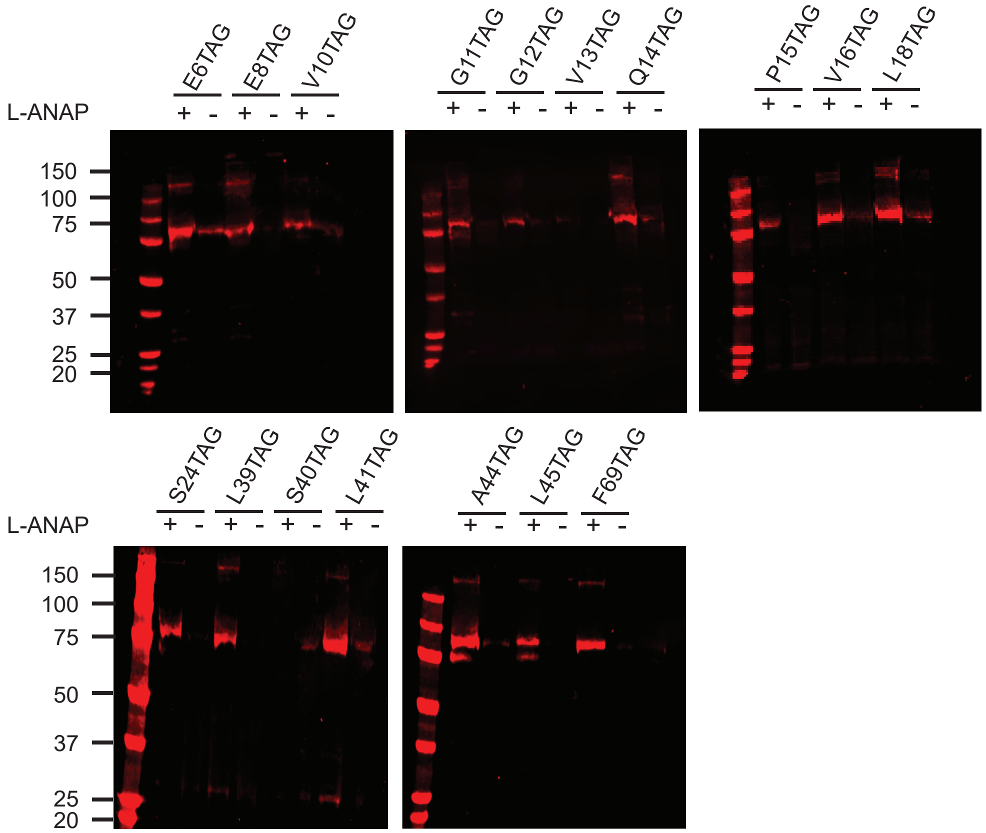


**Figure 2 - figure supplement 1**

Representative western blots for all TAG mutants created in this study. Westerns were blotted with an anti-GFP antibody that recognizes the terminal mCitrine. Each TAG mutant was cultured both with L-ANAP (10μM) and without L-ANAP supplementation. Westerns in this figure were run on Bis-Tris gels, whereas the westerns in Figure 2 were run on Tris-Glycine gels, which results in ASIC running slightly differently between the 2 gel types.

**Figure 2- supplement table 1**

$pH_{50}$ from 2C with N and SEM for each TAG position.

| | | |
|---|---|---|
| C469 WT | $pH_{50}$ | 6.52 |
| | SEM | 0.01 |
| | n | 5 |
| E8TAG | $pH_{50}$ | 6.66 |
| | SEM | 0.01 |
| | n | 3 |
| G11TAG | $pH_{50}$ | 6.58 |
| | SEM | 0.02 |
| | n | 4 |
| Q14TAG | $pH_{50}$ | 6.42 |
| | SEM | 0.02 |
| | n | 5 |
| 505TAG | $pH_{50}$ | 6.58 |
| | SEM | 0.02 |
| | n | 4 |

A

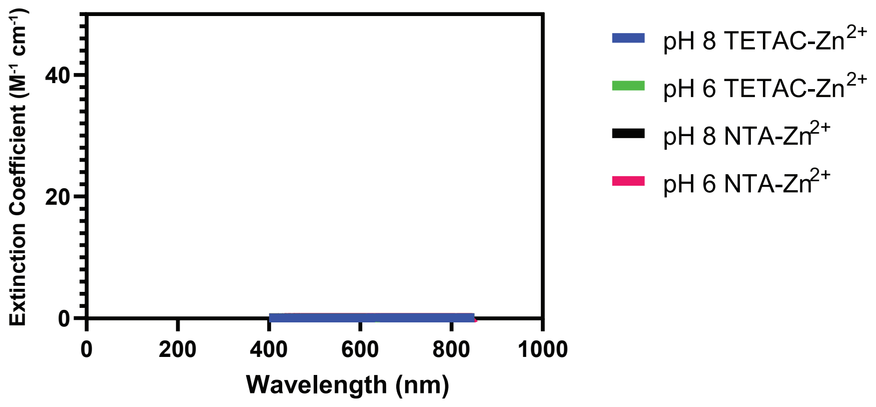


**Figure 3 – figure supplement 1**
(A) Spectral properties of $Zn^{2+}$-TETAC and $Zn^{2+}$-C18-NTA at pH 8 and pH 6 .

**Figure 6- table supplement 1**

P-values of relevant comparisons between NTD TAG positions with cysteine at C469 at pH 8 and pH 6. P values were calculated using two-way ANOVA with Tukey's multiple comparisons test.

| | TAG position comparison | P value |
|---|---|---|
| **pH 8 vs pH 6** | E8TAG pH 8 vs. E8TAG pH 6 | 0.1068 |
| | G11TAG pH 8 vs. G11TAG pH 6 | 0.0091 |
| | Q14TAG pH 8 vs. Q14TAG pH 6 | <0.0001 |
| | S24TAG pH 8 vs. S24TAG pH 6 | 0.0069 |
| | I33TAG pH 8 vs. I33TAG pH 6 | <0.0001 |
| | A44TAG pH 8 vs. A44TAG pH 6 | <0.0001 |
| **pH 8 vs pH 8** | E8TAG pH 8 vs. G11TAG pH 8 | 0.9991 |
| | E8TAG pH 8 vs. Q14TAG pH 8 | 0.0139 |
| | G11TAG pH 8 vs. Q14TAG pH 8 | 0.0071 |
| | S24TAG pH 8 vs. I33TAG pH 8 | 0.0521 |
| | S24TAG pH 8 vs. A44TAG pH 8 | 0.9973 |
| | I33TAG pH 8 vs. A44TAG pH 8 | 0.4091 |
| **pH 6 vs pH 6** | E8TAG pH 6 vs. G11TAG pH 6 | 0.7078 |
| | E8TAG pH 6 vs. Q14TAG pH 6 | <0.0001 |
| | G11TAG pH 6 vs. Q14TAG pH 6 | 0.0818 |
| | S24TAG pH 6 vs. I33TAG pH 6 | 0.9865 |
| | S24TAG pH 6 vs. A44TAG pH 6 | <0.0001 |
| | I33TAG pH 6 vs. A44TAG pH 6 | <0.0001 |

**Figure 6- table supplement 2**

P-values of relevant comparisons between cytosolic NTD TAG positions with cysteine at C477 at pH 8 and pH 6. P values were calculated using two-way ANOVA with Tukey's multiple comparisons test.

| | TAG position comparison | P value |
|---|---|---|
| **pH 8 vs pH 6** | G11TAG pH 8 vs. G11TAG pH 6 | 0.0381 |
| | Q14TAG pH 8 vs. Q14TAG pH 6 | 0.9759 |
| | S24TAG pH 8 vs. S24TAG pH 6 | 0.0075 |
| **pH 8 vs pH 8** | G11TAG pH 8 vs. Q14TAG pH 8 | 0.9732 |
| | G11TAG pH 8 vs. S24TAG pH 8 | 0.0259 |
| | Q14TAG pH 8 vs. S24TAG pH 8 | 0.4234 |
| **pH 6 vs pH 6** | G11TAG pH 6 vs. Q14TAG pH 6 | 0.6374 |
| | G11TAG pH 6 vs. S24TAG pH 6 | 0.2881 |
| | Q14TAG pH 6 vs. S24TAG pH 6 | 0.0015 |

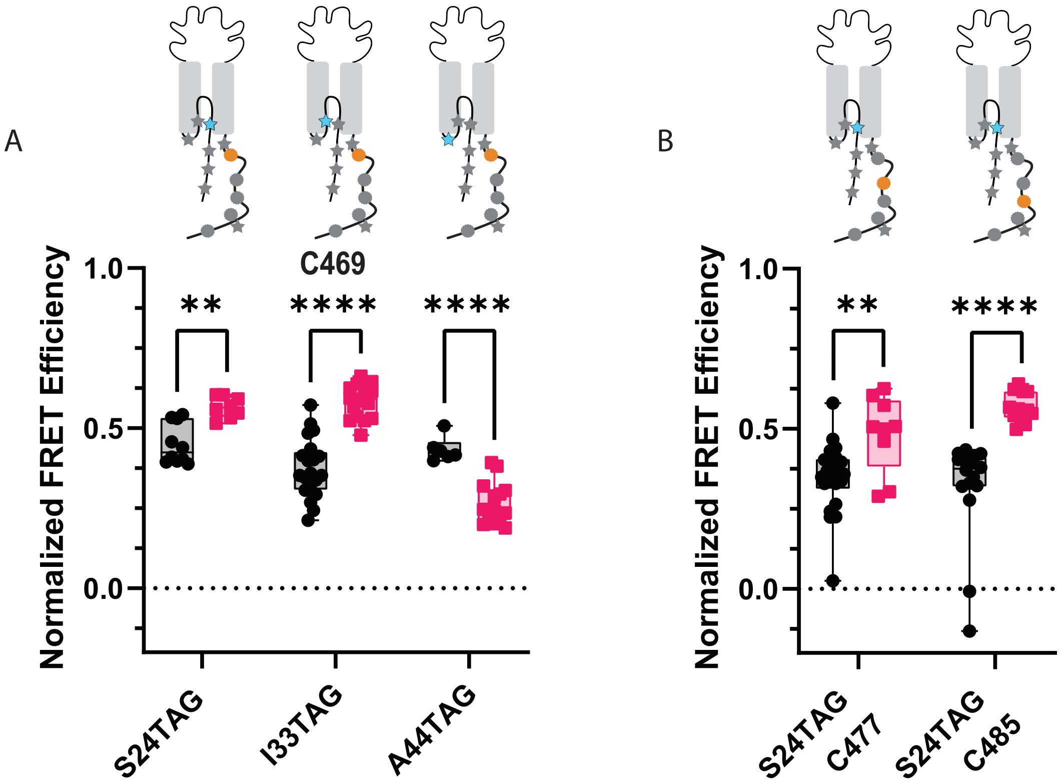


**Figure 7- figure supplement 1**

(A) Normalized FRET efficiency between L-ANAP in the reentrant loop and $Cu^{2+}$-TETAC at position C469 at pH 8 (black) and pH 6 (red). FRET efficiencies, SEM, N and calculated distances are summarized in Table 1. (B) Normalized FRET efficiency between L-ANAP incorporated at S24 and $Cu^{2+}$-TETAC at either C477 or C485. Box and whisker plot with whiskers ranging from minimum to maximum values and the bottom and top edges of the box denoting the 25th and 75th quartiles, respectively. Statistical significance shown using two-way ANOVA with Tukey's multiple comparisons test are denoted between relevant comparison using appropriate asterisks. Ns indicates $p > 0.05$, * indicates $p \le 0.05$, ** indicates $p \le 0.01$, *** indicates $p \le 0.001$, **** indicates $p \le 0.0001$.

**Figure 7- table supplement 1**

P-values of relevant comparisons between cytosolic NTD TAG positions with cysteine at C485 at pH 8 and pH 6. P values were calculated using two-way ANOVA with Tukey's multiple comparisons test.

| | TAG position comparison | P value |
|---|---|---|
| **pH 8 vs pH 6** | E8 TAG pH 8 vs. E8 TAG pH 6 | 0.0065 |
| | G11 TAG pH 8 vs. G11 TAG pH 6 | <0.0001 |
| | Q14 TAG pH 8 vs. Q14 TAG pH 6 | <0.0001 |
| | S24 TAG pH 8 vs. S24 TAG pH 6 | <0.0001 |
| **pH 8 vs pH 8** | E8 TAG pH 8 vs. G11 TAG pH 8 | 0.0005 |
| | E8 TAG pH 8 vs. Q14 TAG pH 8 | 0.0323 |
| | E8 TAG pH 8 vs. S24 TAG pH 8 | <0.0001 |
| | G11 TAG pH 8 vs. Q14 TAG pH 8 | 0.6303 |
| | G11 TAG pH 8 vs. S24 TAG pH 8 | 0.9871 |
| | Q14 TAG pH 8 vs. S24 TAG pH 8 | 0.0833 |
| **pH 6 vs pH 6** | E8 TAG pH 6 vs. G11 TAG pH 6 | <0.0001 |
| | E8 TAG pH 6 vs. Q14 TAG pH 6 | <0.0001 |
| | E8 TAG pH 6 vs. S24 TAG pH 6 | <0.0001 |
| | G11 TAG pH 6 vs. Q14 TAG pH 6 | >0.9999 |

**Figure 7- table supplement 2**

P-values of relevant comparisons between cytosolic NTD TAG positions with cysteine at C505 at pH 8 and pH 6. P values were calculated using two-way ANOVA with Tukey's multiple comparisons test.

| | TAG position comparison | P value |
|---|---|---|
| **pH 8 vs pH 6** | E8 TAG pH 8 vs. E8 TAG pH 6 | 0.0019 |
| | Q14 TAG pH 8 vs. Q14 TAG pH 6 | 0.8454 |
| **pH 8 vs pH 8** | E8 TAG pH 8 vs. Q14 TAG pH 8 | 0.0008 |
| **pH 6 vs pH 6** | E8 TAG pH 6 vs. Q14 TAG pH 6 | 0.0944 |

**Figure 7- table supplement 3**

P-values of relevant comparisons between cytosolic NTD TAG positions with cysteine at C515 at pH 8 and pH 6. P values were calculated using two-way ANOVA with Tukey’s multiple comparisons test.

| | TAG position comparison | P value |
|---|---|---|
| **pH 8 vs pH 6** | E8TAG pH 8 vs. E8TAG pH 6 | <0.0001 |
| | Q14TAG pH 8 vs. Q14TAG pH 6 | <0.0001 |
| **pH 8 vs pH 8** | E8TAG pH 8 vs. Q14TAG pH 8 | <0.0001 |
| **pH 6 vs pH 6** | E8TAG pH 6 vs. Q14TAG pH 6 | <0.0001 |

**Figure 8- table supplement 1**

P-values of relevant comparisons between NTD TAG positions with the plasma membrane at pH 8 and pH 6. P values were calculated using two-way ANOVA with Tukey's multiple comparisons test.

| | TAG position comparison | P value |
|---|---|---|
| **pH 8 vs pH 6** | E8TAG pH 8 vs. E8TAG pH 6 | 0.9998 |
| | G11TAG pH 8 vs. G11TAG pH 6 | 0.9975 |
| | Q14TAG pH 8 vs. Q14TAG pH 6 | 0.0032 |
| | S24TAG pH 8 vs. S24TAG pH 6 | >0.9999 |
| | I33TAG pH 8 vs. I33TAG pH 6 | 0.0001 |
| **pH 8 vs pH 8** | E8TAG pH 8 vs. G11TAG pH 8 | 0.9949 |
| | E8TAG pH 8 vs. Q14TAG pH 8 | >0.9999 |
| | G11TAG pH 8 vs. Q14TAG pH 8 | >0.9999 |
| | S24TAG pH 8 vs. I33TAG pH 8 | 0.0092 |
| **pH 6 vs pH 6** | E8TAG pH 6 vs. G11TAG pH 6 | 0.1550 |
| | E8TAG pH 6 vs. Q14TAG pH 6 | <0.0001 |
| | G11TAG pH 6 vs. Q14TAG pH 6 | 0.0576 |
| | S24TAG pH 6 vs. I33TAG pH 6 | >0.9999 |

**Figure 8- table supplement 2**

P-values of relevant comparisons between CTD TAG positions with the plasma membrane at pH 8 and pH 6. P values were calculated using two-way ANOVA with Tukey's multiple comparisons test.

| | TAG position comparison | P value |
|---|---|---|
| **pH 8 vs pH 6** | 464TAG pH 8 vs. 464TAG pH 6 | 0.8153 |
| | 505TAG pH 8 vs. 505TAG pH 6 | <0.0001 |
| **pH 8 vs pH 8** | 464TAG pH 8 vs. 505TAG pH 8 | 0.8693 |
| **pH 6 vs pH 6** | 464TAG pH 6 vs. 505TAG pH 6 | 0.0366 |
